# Model-based fMRI Reveals Dissimilarity Processes Underlying Base Rate Neglect

**DOI:** 10.1101/130716

**Authors:** Sean R. O’Bryan, Darrell A. Worthy, Evan J. Livesey, Tyler Davis

**Author notes:** CONTACT: Correspondence to: Sean R. O’Bryan Texas Tech University, Department of Psychological Sciences MS 2051, Lubbock, TX 79409.

## Abstract

Extensive evidence suggests that people use base rate information inconsistently in decision making. A classic example is the inverse base rate effect (IBRE), whereby participants classify ambiguous stimuli sharing features of both common and rare categories as members of the rare category. Computational models of the IBRE have either posited that it arises from associative similarity-based mechanisms or dissimilarity-based processes that may depend upon higher-level inference. Here we develop a hybrid model, which posits that similarity- and dissimilarity-based evidence both contribute to the IBRE, and test it using functional magnetic resonance imaging data collected from human subjects completing an IBRE task. Consistent with our model, multivoxel pattern analysis reveals that activation patterns on ambiguous test trials contain information consistent with dissimilarity-based processing. Further, trial-by-trial activation in left rostrolateral prefrontal cortex tracks model-based predictions for dissimilarity-based processing, consistent with theories positing a role for higher-level symbolic processing in the IBRE.

## Introduction

Does this patient have influenza or Ebola virus? Categorization is a fundamental process that underlies many important decisions. Categories, such as viruses, often have different relative frequencies or base rates. Influenza, for example, is very common and infects millions of people worldwide each year, whereas Ebola virus tends to have infection rates that are orders of magnitude lower.

One critical question is how people use such base rate information when making categorization decisions. Research so far has suggested that people tend to be, at best, inconsistent in their use of base rate information. Both in realistic studies with medical professionals and artificial categorization tasks in the lab, when confronted with examples that share characteristics with both rare and common categories, people show a tendency to predict the rare category much more often than the base rates would suggest (Tversky & Kahneman, 1974; Casscells, Schoenberger & Graboy, 1978; Bravata, 2000). In an extreme case, known as the inverse base rate effect (IBRE), people may even predict rare categories as more likely than common ones (Medin & Edelson, 1988). For example, in an IBRE context, a patient presenting with cough (a characteristic feature of influenza) and unexplained bleeding (a characteristic feature of Ebola), may be more likely to be diagnosed with Ebola than influenza.

The mechanisms that lead to base rate neglect are currently undetermined at both the cognitive and neural levels. Computationally, according to influential work with similarity-based categorization models (Medin & Edelson, 1988; Kruschke, 1996, 2001), the IBRE arises from differential selective attention to features for common and rare categories. Specifically, participants learn to attend more strongly to features of rare categories, making ambiguous cases seem more similar to rare categories and thus more likely to be rare category members. In terms of the flu example, participants may attend more to the unexplained bleeding feature of the rarer Ebola virus category, and thus predict Ebola when confronted with a patient with both features.

Similarity-based category learning models have strong support in the neurobiological category learning literature. Model-based predictions for how similar items are to stored category representations have been shown to correlate with activation in the medial temporal lobes (MTL; Davis, Love & Preston, 2012a; 2012b). Moreover, at a finer-grained level, multivoxel activation patterns in the MTL have been shown to contain information associated with higher-order similarity relationships between category members anticipated by similarity-based models (Davis & Poldrack, 2014), including those predicted by differences in selective attention (Mack, Love & Preston, 2016). The dorsolateral prefrontal cortex (dlPFC) tends to track predictions of choice uncertainty from similarity-based models, whereas ventromedial PFC (vmPFC) tends to track estimates of high choice accuracy or model-confidence (Davis, Goldwater & Giron, 2017).

Despite the strong cognitive and neural evidence for similarity-based models, it remains an open question whether they provide a complete account of IBRE-like phenomena. One alternative that has been proposed is that people’s choice of rare categories when confronted with conflicting information may stem from the reliance on dissimilarity processes, either on their own, or in addition to similarity-based processes. According to theories that focus on dissimilarity-based processes, people build strong expectations of the common category; thus they view items containing features inconsistent with these expectations as more likely to be members of the rare category (Juslin, Wennerholm & Winman, 2001; Winman et al., 2005). For example, a doctor may have seen thousands of cases of flu, none with unexplained bleeding, and thus rule out influenza and choose Ebola virus based on these expectations. In these cases, it is *dissimilarity* to members of the common category that drives choice, rather than the similarity to rare category members per se.

Formal models positing dissimilarity processes have so far been explicitly dual-process oriented. For example, ELMO, a computational model that incorporates a choice elimination decision based on dissimilarity argues that such elimination depends on explicit reasoning processes that are separate from similarity-based processes that arise on other trials (Juslin, Wennerholm & Winman, 2001). In the present study, we propose a new account based on a recently proposed dissimilarity-based extension of the generalized context model, the dissGCM (Stewart & Morin, 2007). This account uses the exact same basic similarity computations as standard similarity-based models (e.g., Nosofsky, 1986), but allows similarities and dissimilarities to stored exemplars to be used as evidence for a category. In terms of the above example, dissimilarity to influenza can be used as evidence for Ebola (and vice versa).

As specified computationally, the dissGCM is agnostic about whether dissimilarity-based evidence constitutes a different cognitive or neurobiological mechanism from similarity-based evidence. On one hand, the dissGCM has no fundamentally different computations from a basic similarity process; as detailed below, dissimilarity is a simple transformation of similarity. On the other hand, it is possible that dissimilarity processes require manipulating similarity relationships between category representations in a more symbolic or abstract manner, as anticipated by previous dissimilarity theories.

Higher-level cognitive control mechanisms are thought to depend upon a hierarchy of abstraction in the lateral PFC along the rostral-caudal axis (Badre & D’Esposito, 2007, 2009). At the apex of this hierarchy is the rostrolateral PFC (rlPFC), a region often implicated in tasks that require people to generalize across abstract, symbolic representations. For example, relational reasoning tasks like Raven’s progressive matrices and rule-based tasks involving abstract relations are thought to depend on left rlPFC (Christroff et al., 2001; Bunge et al., 2005, 2009; Davis, Goldwater, & Giron, 2017). In addition to its role in generalizing abstract, relational rules, we have recently found left rlPFC to be involved in rule evaluation and novel generalization processes for simpler feature-based rules in categorization tasks (Paniukov & Davis, 2018). In the present study, dissimilarity-based generalization to novel feature pairings may depend on rule evaluation processes in the rlPFC more so than simple similarity-based processing, if studies anticipating that dissimilarity-based processes depend more upon higher-level symbolic rules are correct (Juslin, Wennerholm, & Winman, 2001; Winman et al., 2005).

Here we test the dissGCM by incorporating its predictions into an analysis of fMRI data collected from participants completing a standard IBRE task (Medin & Edelson, 1988; Kruschke, 1996). We first examine whether activation patterns elicited during conflicting trials in the IBRE task are consistent with participants thinking more about the rare category, as predicted by pure similarity-based accounts, or thinking more about (dissimilarity to) the common category, as predicted by the dissGCM. To this end, we use representational similarity analysis (RSA; Kriegeskorte et al., 2008) to decode which features of the stimuli are most strongly activated while participants are categorizing the conflicting items. This analysis is based on recent work in the broader memory literature establishing that it is possible to decode whether participants are thinking about or retrieving particular object categories from memory based on their activation patterns in ventral temporal cortex (Rissman & Wagner, 2012; Haxby, Connolly, & Guntupalli, 2014). Here we use the real world visual categories faces, scenes, and objects as stimulus features. These visual categories have a well-defined representational topography across the cortex (Haxby et al., 2001; Grill-Spector & Weiner, 2014), allowing us to predict whether participants are thinking about a particular stimulus feature (faces, scenes, or objects) by computing similarities between activation patterns elicited for the key IBRE trials and feature-specific patterns from an independent localizer scan. By crossing the visual stimulus features with our category structure (Figure 1), we create situations where a rare category is associated with one feature type (e.g., a scene) whereas a common category is associated with another feature type (e.g., an object). The extent to which each type of information is active can then be compared to determine whether participants are thinking more about common or rare categories on a trial, and thus answer whether their neural activation patterns are more consistent with pure similarity or dissGCM’s combined dissimilarity and similarity processes.

**Figure 1.**
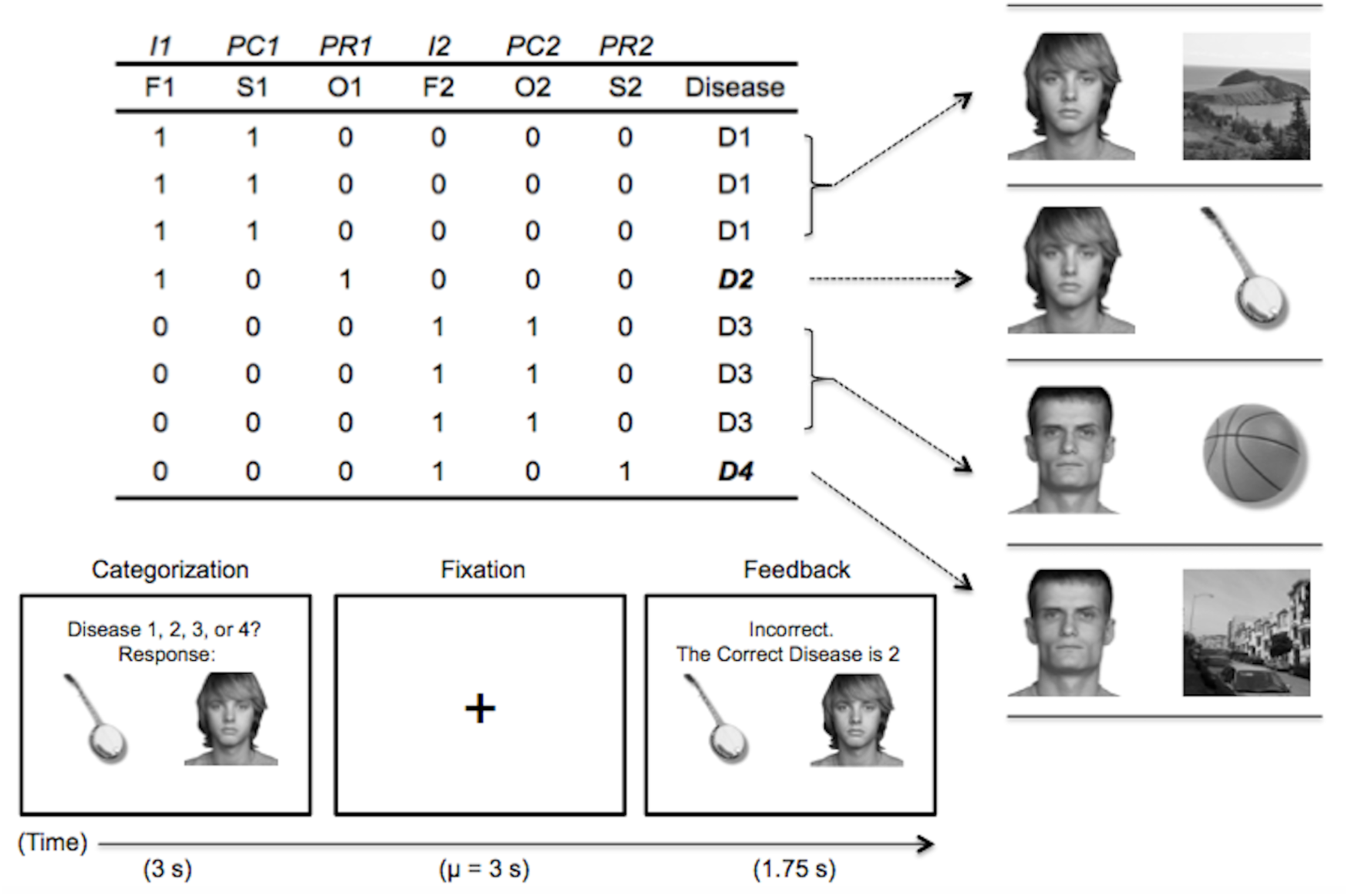
Abstract task design and an example trial. In the headings, I = imperfect predictor, PC = common perfect predictor, PR = rare perfect predictor. The second row refers to the visual category used for each stimulus feature: F = face, S = scene, O = object. Each following row corresponds to a learning trial, with a “1” indicating the presence of the feature and “0” indicating its absence.

In addition to our multivoxel analysis, we also test whether the dissimilarity-based evidence may tap distinct brain regions, such as the rlPFC, beyond those involved with similarity-based evidence. To this end, we take trial-by-trial predictions for how much similarity-and dissimilarity-based evidence contribute to the winning category choice and use these predictions as regressors in fMRI analysis. We anticipated that the MTL and vmPFC would be positively associated with similarity-based evidence, whereas dlPFC would be negatively associated with similarity-based evidence. Contrastingly, we expected rlPFC to track estimates of dissimilarity-based evidence.

## Model

The dissimilarity generalized context model (dissGCM) is based off of the original generalized context model (Nosofsky, 1986), but allows for dissimilarity to be used as evidence for a decision (Stewart & Morin, 2007). The model posits that people represent stimuli as points in a multidimensional feature space. Categorization judgments are based off of distances between probe stimuli *Si* and stored exemplars *Sj* along each dimension *k*:

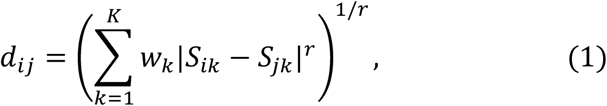

where *r* defines the metric of the space, here assumed to be one (city-block). The *w_k_* indicates dimensional attention weights, which have the function of stretching the distance along strongly attended dimensions, and are constrained to sum to one.

Distances are converted to similarities via an exponential transform:

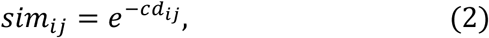

where *c* is a specificity parameter that controls the rate at which similarity decays as a function of distance.

Like the standard GCM, similarities to all exemplars of each category are summed into evidence for each category. However, in the dissGCM, evidence that an item is dissimilar to other categories is also used as evidence for a category. For example, evidence for Disease 1 includes not only an item’s similarity to members of Disease 1, but also its dissimilarity to other diseases. The overall evidence, *v*, for a category *C_A_*, given stimulus *S_i_* is:

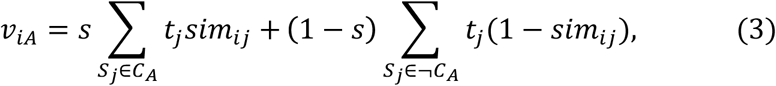

where *s* is a free parameter that determines how much the model weights similarity versus dissimilarity. The parameter *t_j_* reflects exemplar-specific memory strength, which we fix at each exemplar’s true base rate during learning (1 for rare category exemplars, 3 for common category exemplars). Here, we also make the assumption that exemplars only contribute evidence (similarity or dissimilarity) if they have at least one positive feature match with a probe stimulus.

The model makes a prediction for how likely an item is to be classified as a member of a given category *C_A_* by:

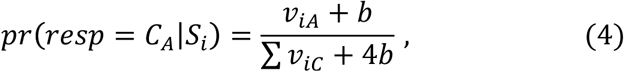

where *b* is a free parameter that reflects the baseline level of similarity for a category that has zero positive feature matches. More generally, this parameter ensures that no predicted probabilities are zero or one, which interferes with the maximum likelihood-based model fits.

The model was fit to the group response frequencies for each option by minimizing the −2 * Log Likelihood (G^2^) using a differential evolution function optimizer. The overall fit was 4,314.588. The best fitting parameters for each of the dimension weights were *w*_1_ (face 1) = 0.277, *w*_2_ (common scene) = 0.665, *w*_3_ (rare object) = 0.887, *w*_4_ (face 2) = 0.170, *w*_5_ (common object) = 0.712, and *w*_6_ (rare scene) = 0.879); *c* = 9.05; *s* = 0.946; *b* = 0.023.

For the model-based neuroimaging analysis, we break the evidence *v* for the winning (most probable) category into separate measures indicating the similarity-based evidence (the summed similarity to the winning category) and dissimilarity-based evidence (summed dissimilarity to other categories). Likewise, for the multivoxel analysis we examine how much each category’s exemplars (common and rare) contribute to the rare response for ambiguous items to predict how strongly participants should be activating information associated with each category.

## Results

### Behavioral Results

Learning curves over the 12 learning blocks for common and rare disease item pairs are shown in Figure 2. All subjects reached greater than 90% accuracy over the last 4 blocks (*M* = 98.1%, *SD* = 2.4%, range = 93.5 – 100%). Mean choice performance in the first block was above chance (25%) for both common (*M* = 63.6%) and rare (*M* = 43.2%) feature pairs. Consistent with previous IBRE studies, a mixed effects ANOVA revealed a significant block by feature type interaction, *F* (1, 21) = 9.87, *p* = .005: the common diseases were learned more quickly than the rare diseases, with prediction accuracy for the common and rare feature pairs becoming comparable by the fifth learning block.

**Figure 2.**
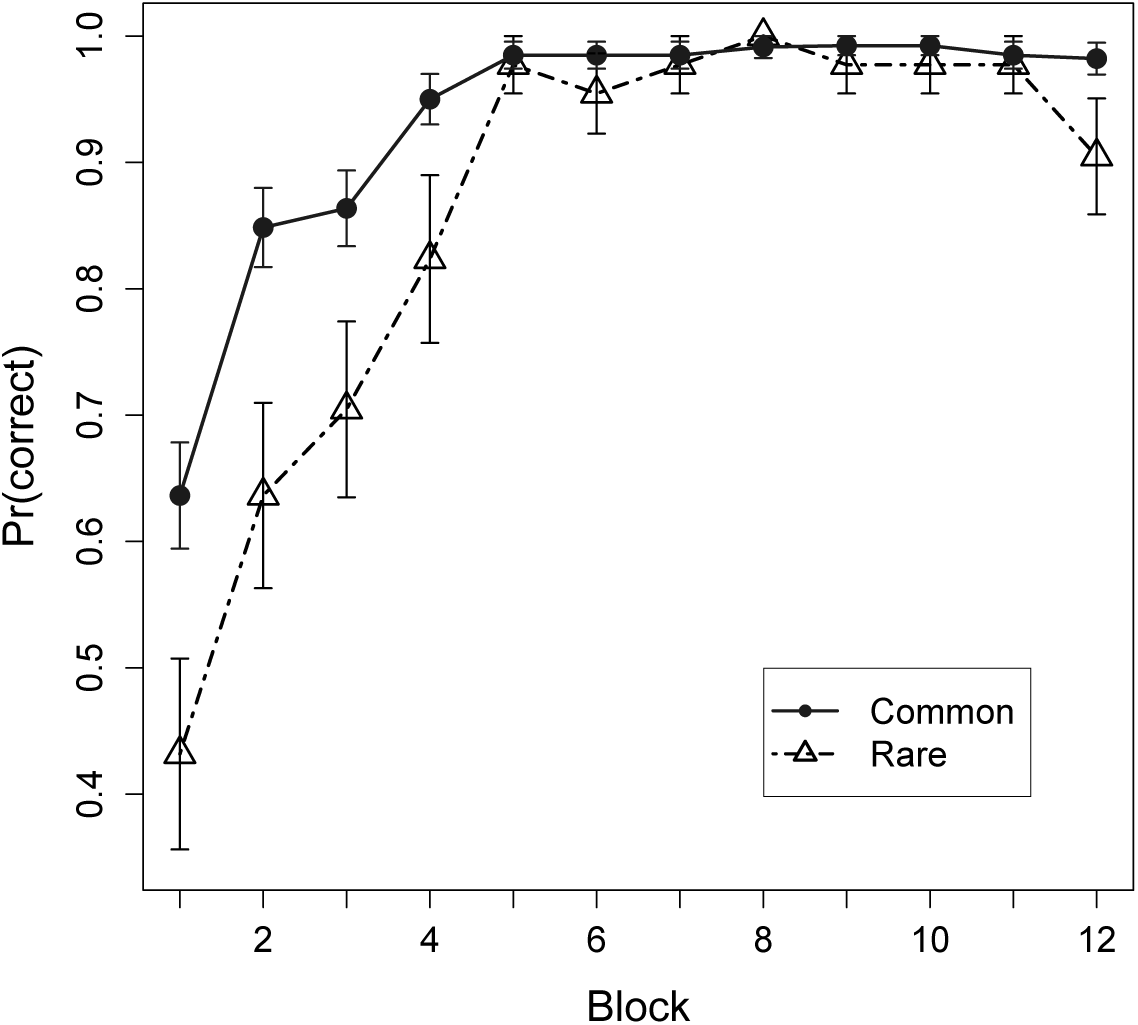
Learning curves. Points depict proportions correct for common and rare disease predictions over the 12 blocks of the training phase (mean ± SEM).

Choice probabilities and dissGCM-derived predictions for each of the test items are summarized in Table 1. Consistent with an inverse base-rate effect, participants were numerically more likely to classify ambiguous test stimuli (combinations of rare and common features) as members of the relevant rare category (*M* = 47.6%) than the relevant common category (*M* = 43.4%). A one-sample t-test revealed that the percentage of rare responding on ambiguous trials was significantly higher than the 1/4 base rate for the rare category, *t* (21) = 8.11, *p* < .001.

**Table 1.**
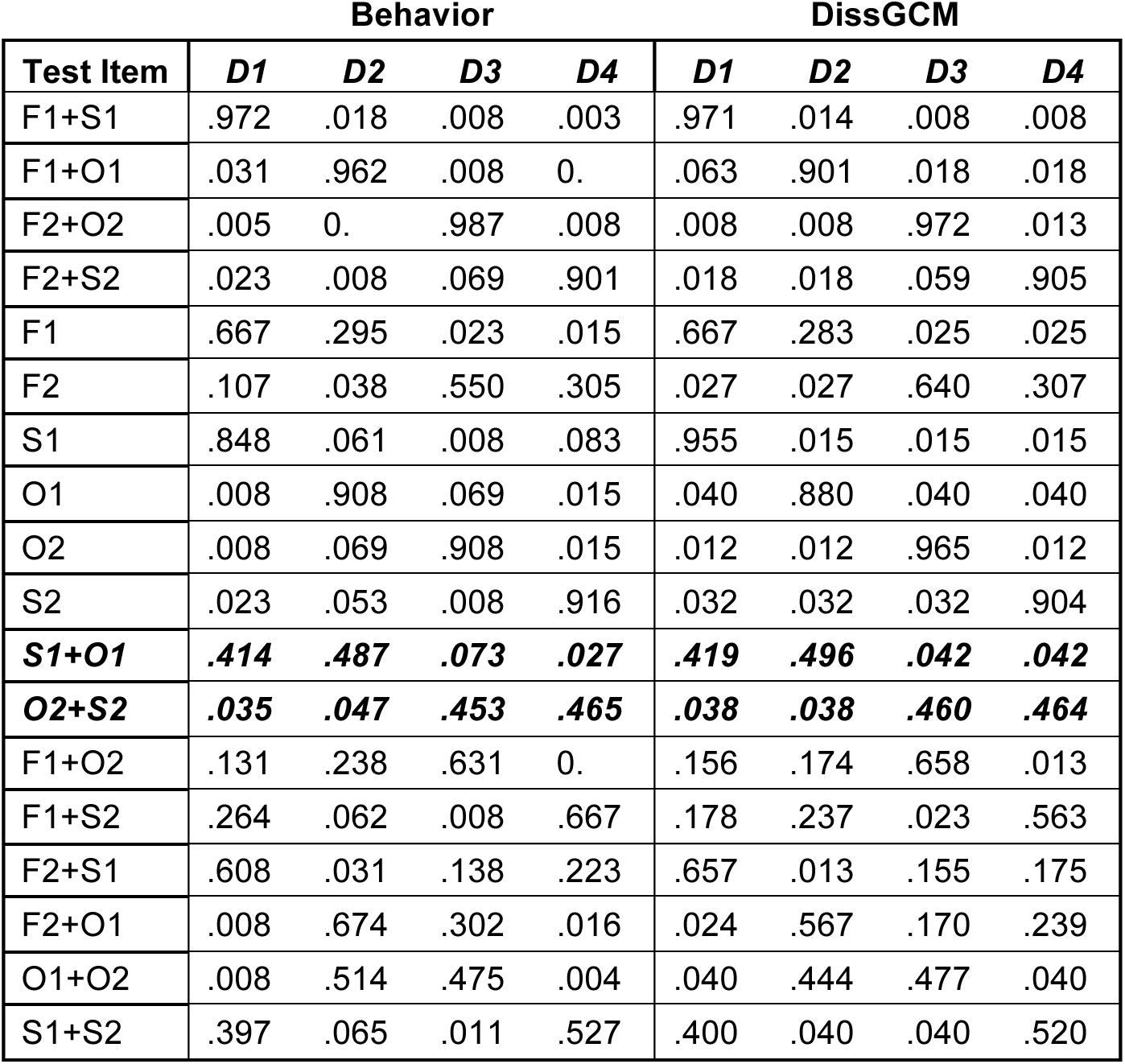
Observed and dissGCM-predicted response probabilities for the test phase. The feature combinations presented at test are listed in the leftmost column: F = face, S = scene, O = object. In the headings, D1 – D4 correspond to the four possible category responses (diseases). Bold, italicized values indicate results for the key ambigous stimuli in which a scene was paired with an object.

| Test Item | Behavior |  |  |  | DissGCM |  |  |  |
| --- | --- | --- | --- | --- | --- | --- | --- | --- |
|  | D1 | D2 | D3 | D4 | D1 | D2 | D3 | D4 |
| F1+S1 | .972 | .018 | .008 | .003 | .971 | .014 | .008 | .008 |
| F1+O1 | .031 | .962 | .008 | 0. | .063 | .901 | .018 | .018 |
| F2+O2 | .005 | 0. | .987 | .008 | .008 | .008 | .972 | .013 |
| F2+S2 | .023 | .008 | .069 | .901 | .018 | .018 | .059 | .905 |
| F1 | .667 | .295 | .023 | .015 | .667 | .283 | .025 | .025 |
| F2 | .107 | .038 | .550 | .305 | .027 | .027 | .640 | .307 |
| S1 | .848 | .061 | .008 | .083 | .955 | .015 | .015 | .015 |
| O1 | .008 | .908 | .069 | .015 | .040 | .880 | .040 | .040 |
| O2 | .008 | .069 | .908 | .015 | .012 | .012 | .965 | .012 |
| S2 | .023 | .053 | .008 | .916 | .032 | .032 | .032 | .904 |
| **S1+O1** | **.414** | **.487** | **.073** | **.027** | **.419** | **.496** | **.042** | **.042** |
| **O2+S2** | **.035** | **.047** | **.453** | **.465** | **.038** | **.038** | **.460** | **.464** |
| F1+O2 | .131 | .238 | .631 | 0. | .156 | .174 | .658 | .013 |
| F1+S2 | .264 | .062 | .008 | .667 | .178 | .237 | .023 | .563 |
| F2+S1 | .608 | .031 | .138 | .223 | .657 | .013 | .155 | .175 |
| F2+O1 | .008 | .674 | .302 | .016 | .024 | .567 | .170 | .239 |
| O1+O2 | .008 | .514 | .475 | .004 | .040 | .444 | .477 | .040 |
| S1+S2 | .397 | .065 | .011 | .527 | .400 | .040 | .040 | .520 |

### Multivoxel Results

#### Test Phase

The primary goal of the multivoxel analysis was to decode, for the ambiguous stimuli, whether participants were thinking more about the common or rare category when they make the choice to classify the stimulus as rare. Specifically, for the bold italicized stimuli listed in Table 1, we tested whether participants’ activation patterns were more similar to localizer activation patterns associated with scenes when a scene was the common feature (and object was rare) and more similar to those of objects when an object was the common feature (and scene was rare).

The prediction that information associated with the common category should be more active on ambiguous trials is derived from the dissGCM. The model posits that the higher probability for a rare response is based on the contribution that dissimilarity to the common category makes to the evidence for the rare category. Indeed, in the fitted version of the model, the evidence for rare contributed by similarity to the rare category exemplar was nearly half the evidence contributed by dissimilarity to the common category exemplar (rare = 0.088; common = 0.153, in the dissGCM’s attention weighted similarity units).

Neural similarities to both visual stimulus categories on the ambiguous test trials are depicted in Figure 3. Consistent with the dissGCM’s predictions, a linear mixed effects model revealed that when participants made a rare choice, their activation patterns were most similar to whichever visual stimulus category (scenes or objects) was associated with the common category, *t* (21) = 2.78, *p* = .011. Interestingly, there was no significant difference between pattern similarity for rare and common features when participants made a common response, *t* (21) = 0.45, *p* =.653. This pattern, whereby participants tended to only differentially activate patterns associated with common features when they made a rare response, manifested in a significant interaction, *t* (42) = −2.22, *p* = .032.

**Figure 3.**
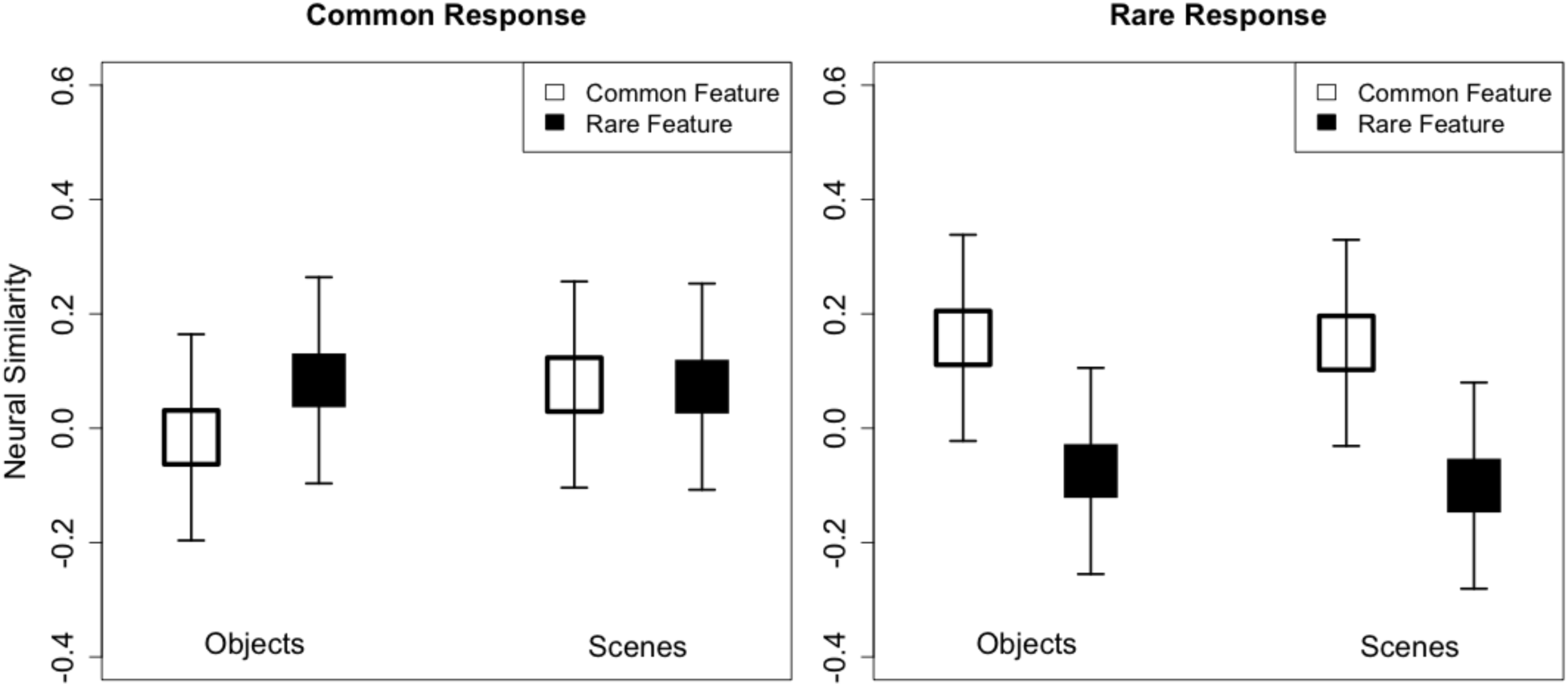
Neural similarity to visual stimulus categories for ambiguous trials in which participants made common (left) and rare (right) responses (mean ± SEM). The white squares indicate similarity for trials in which an object or scene was the common category feature. Black squares indicate similarity for trials in which an object or scene was the rare category feature.

#### Learning Phase

Beyond our primary questions about test phase activation, multivoxel analysis of the learning phase can provide additional information about how participants were processing stimuli in the present task. Generally, both similarity-based models and dissimilarity-based models like the dissGCM predict that features which are most informative about the correct category will contribute more to categorization decisions during learning. With respect to multivoxel predictions, this means that activation patterns elicited during learning should contain more information about the predictive features (objects or scenes) than non-predictive features (faces), and both of these types of information should be activated more strongly than non-present features. Figure 4 depicts mean pattern similarities for predictive, non-predictive, and non-present visual stimulus categories during the learning phase for both common and rare disease trials. As anticipated, a one-way within-subjects ANOVA collapsed across trial type revealed that neural similarity to the visual category was the strongest for perfectly predictive features (*M =* .065), followed by the non-predictive but present features (*M* = -.050) and the non-present features (*M* = -.145), *F* (2, 42) = 41.0, *p* < .001. This finding, whereby activation patterns elicited for stimuli during learning are most similar to predictive features, is consistent with recent studies using MVPA to measure dimensional selective attention in categorization and reinforcement learning (Mack et al., 2013; 2016; Leong et al., 2017; O’Bryan et al., 2018).

**Figure 4.**
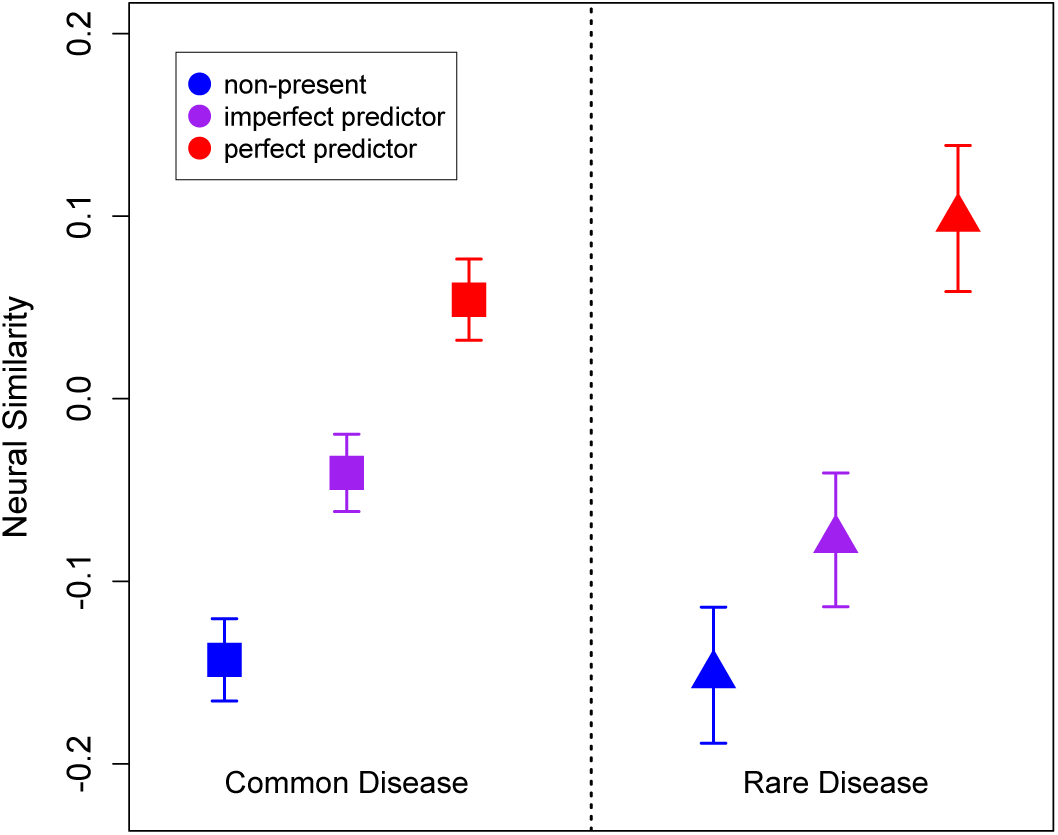
Multivoxel pattern similarity to each feature type during the learning phase (mean ± SEM). The left panel is for trials predictive of a common disease, and the right for trials predictive of a rare disease. Red points represent perfect predictors, purple points represent imperfect predictors (faces), and blue points represent non-present features.

Although the greater contribution of predictive features to multivoxel activation patterns during learning is a straightforward prediction that is consistent with any model, a further question is how such activation patterns during learning contribute to later test performance. As with the test phase, dissimilarity-based theories make the somewhat counterintuitive prediction that it is specifically what people learn about the common category that is driving later choices of the rare category. One way to test this using activation patterns elicited during learning is to examine how the activation of rare and common information patterns is associated with the extent to which individual participants exhibit the inverse base rate effect later during the test phase. Consistent with dissimilarity-based accounts, we found that greater activation of neural patterns associated with the common features during learning was correlated with a higher proportion of rare choices on the ambiguous test trials, *r* = .550, *t* (20) = 2.95, *p* = .008. Alternatively, we found no relationship between activation of neural patterns associated with the rare features during learning and choice proportions on IBRE trials, *r* = .158, *t* (20) = .713, *p* = .483. Figure 5 depicts the associations between neural similarity to common and rare features during learning and base-rate neglect.

**Figure 5.**
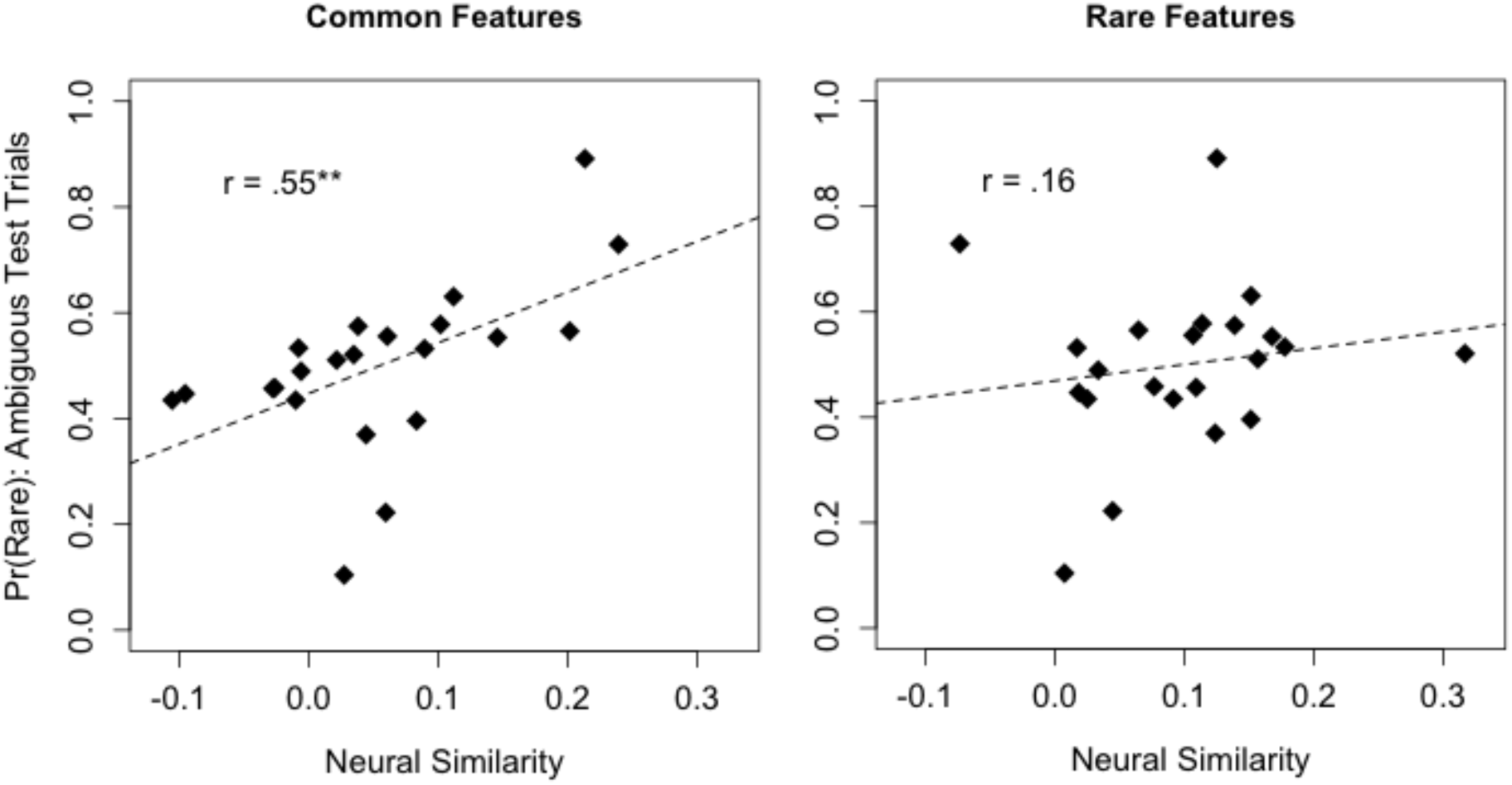
Associations between multivoxel pattern similarity to common (left) and rare (right) features during the learning phase and individual differences in base-rate neglect. The y-axis represents the proportion of rare responses made by each subject on ambiguous test trials. ‘**’ = *p* < 0.01.

### Model-based Univariate Results

By revealing a link between activation of common feature patterns and the IBRE, our multivoxel results suggest that dissimilarity-based evidence contributes to choice behavior in the present task. However, it remains an open question whether such dissimilarity processes involve distinct neural or cognitive mechanisms beyond those thought to underlie basic similarity processes.

To test whether similarity-and dissimilarity-based evidence rely on different brain regions, we modeled univariate voxel-wise activation using trial-by-trial estimates of similarity-and dissimilarity-based evidence derived from the dissGCM (Figure 6). We found activation in the MTL and vmPFC that was positively correlated with similarity-based evidence, whereas the dlPFC and posterior parietal cortex were negatively correlated with similarity-based evidence. These results are consistent with findings from other model-based fMRI studies suggesting that the MTL is involved in similarity-based retrieval (Davis, Love & Preston, 2012a, 2012b), and that the vmPFC and dlPFC track higher-confidence or higher-uncertainty categorization decisions, respectively (DeGutis & D’Esposito, 2007; Seger et al., 2015; Davis, Goldwater & Giron, 2017; O’Bryan et al., 2018). Contrastingly, dissimilarity-based evidence was positively correlated with activation in the left rlPFC, consistent with our hypothesis that this type of evidence might encourage more symbolic processes believed to underlie the rlPFC’s contribution to category learning (Davis, Goldwater & Giron, 2017; Paniukov & Davis, 2018). No clusters were significantly negatively associated with dissimilarity-based evidence.

**Figure 6.**
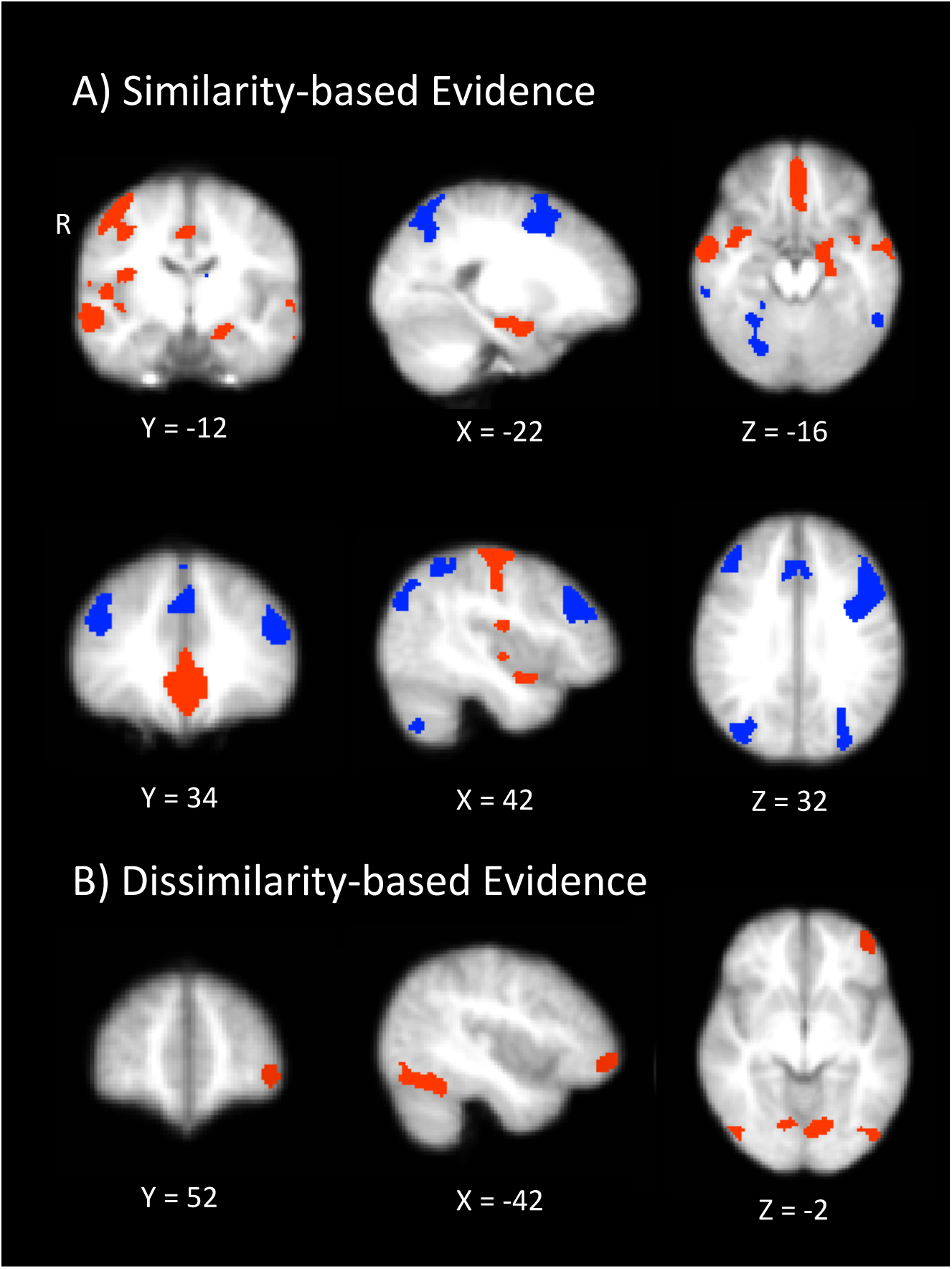
Results from the model-based univariate analysis. A) Depicts activation that tracks similarity-based contributions to choice. Red depicts activation positively correlated with similarity-based contributions whereas blue depicts negatively correlated activation. B) Depicts brain regions that are positively correlated with dissimilarity-based contributions to choice

**Table 2.** Activated clusters and peaks for the fMRI results in Figure 6.

| Contrast | Regions | Peak<br>*t*-value | Peak MNI<br>Coordinates (x,y,z) | Number of<br>voxels | Cluster *p* |
| --- | --- | --- | --- | --- | --- |
| Similarity > 0 |  |  |  |  |  |
| | Rostral and ventral medial prefrontal cortex | 7.76 | 0, 54, -2 | 2268 | $p < .001$ |
| | Middle temporal gyrus | 5.74 | 64, -6, -14 | 1050 | $p = .016$ |
| | Precentral gyrus | 7.65 | 42, -16, 62 | 1031 | $p < .001$ |
| | Precentral gyrus | 4.81 | 0, -30, 58 | 463 | $p = .008$ |
| | Middle temporal gyrus | 4.98 | -62, -2, -16 | 322 | $p = .016$ |
| | Parietal operculum cortex | 5.74 | -34, -30, 18 | 282 | $p = .027$ |
| | Hippocampus | 5.25 | -22, -18, -16 | 271 | $p = .019$ |
| | Lateral occipital cortex (inferior) | 4.68 | 50, -72, 12 | 157 | $p = .039$ |
| Similarity < 0 |  |  |  |  |  |
| | Superior Parietal & Lateral occipital cortex (superior) | 8.43 | -44, -46, 58 | 5123 | $p < .001$ |
| | Middle frontal gyrus | 7.01 | -52, 12, 36 | 2491 | $p < .001$ |
| | Dorsal medial PFC | 9.00 | -2, 18, 46 | 917 | $p < .001$ |
| | Cerebellum | 6.11 | 28, -64, -28 | 769 | $p = .002$ |
| | Middle frontal gyrus | 6.23 | 32, 2, 62 | 730 | $p = .002$ |
| | Middle frontal gyrus | 5.89 | 44, 36, 30 | 391 | $p = .009$ |
| | Inferior temporal gyrus | 6.09 | -54, -52, -12 | 375 | $p = .009$ |
| | Inferior temporal gyrus | 6.53 | 60, -52, -10 | 271 | $p = .016$ |
| | Cerebellum | 6.08 | -28, -60, -32 | 186 | $p = .031$ |
| | Thalamus | 4.88 | -10, -18, 10 | 175 | $p = .038$ |
| Dissimilarity > 0 |  |  |  |  |  |
| | Occipital cortex | 7.55 | 12, -78, 12 | 1372 | $p < .001$ |
| | Fusiform and Lateral occipital cortex (inferior) | 6.82 | 42, -60, -18 | 865 | $p = .002$ |
| | Fusiform and Lateral occipital cortex (inferior) | 5.97 | -36, -52, -18 | 575 | $p = .003$ |
| | Middle frontal gyrus | 5.00 | 54, 16, 34 | 255 | $p = .020$ |
| | Frontal Pole (rostrolateral PFC) | 5.93 | -42, 52, -6 | 130 | $p = .047$ |

## Discussion

The present study employed model-based fMRI to test how similarity and dissimilarity contribute to the inverse base rate effect (IBRE) and how these types of evidence relate to neural mechanisms that support category learning. The dominant theory behind the IBRE suggests that it arises from attentional processes that make ambiguous items containing features of rare and common categories seem more similar to members of the rare category. Here we find support for the hypothesis that dissimilarity-based evidence also contributes to the IBRE: people may categorize the ambiguous stimuli as members of the rare category not only because of their similarity to the rare category, but also because of their dissimilarity to members of the common category.

The dissGCM, an extension of the GCM that allows for the use of dissimilarity-based evidence in categorization behavior, predicted two novel observations in the neuroimaging data. First, as predicted by the dissGCM’s relative contribution of similarity-and dissimilarity-based evidence during the ambiguous trials, multivoxel analysis suggested stronger activation of patterns associated with features of the common category when participants classified ambiguous stimuli as rare. Second, model-based univariate analysis revealed that measures of similarity-and dissimilarity-based evidence had unique neural topographies. Similarity-based evidence was positively correlated with regions of the hippocampus and vmPFC and negatively correlated with dlPFC. Dissimilarity-based evidence was positively correlated with the left rlPFC.

The present results raise several important questions about the cognitive and neural mechanisms underlying people’s use of base rate information. Previous theories arguing for dissimilarity-like processes as explanations of IBRE have argued that they arise from mechanisms rooted in higher-level propositional logic that fundamentally differ from the similarity-based mechanisms posited by dominant theories (Juslin, Wennerholm & Winman, 2001). As illustrated by the dissGCM, such dissimilarity-based processes can be viewed as simple extensions of similarity-based processing and need not depend on the existence of a functionally separate categorization system. At the same time, our neuroimaging results suggest dissimilarity, but not similarity-based evidence may arise from processing in rlPFC regions that are known to be involved with higher-level reasoning and problem solving (Christoff et al., 2001; Bunge et al., 2005, 2009). One possibility for reconciling these theories is that the dissimilarity-based evidence involves more abstract or symbolic feature processing than pure similarity processes, and this additional processing taps rlPFC regions. This is consistent with our recent model-based fMRI results which demonstrate that rlPFC tracks measures of relational encoding in category learning, but otherwise this type of category learning may rely on the same basic similarity-based mechanisms as simpler feature-based learning (Davis, Goldwater, & Giron, 2017).

By establishing that the rlPFC is engaged when participants incorporate dissimilarity-based evidence into categorization decisions, our research adds to a growing literature aiming to pinpoint a domain-general computational role for this region. A common thread among tasks shown to engage the rlPFC is that they tend to involve combining across disparate representations to form the basis for a decision – whether those representations are comprised of confidence estimates and subjective value (De Martino et al., 2013), visual features and their relations (Bunge et al., 2004, 2009; Davis, Goldwater, & Giron, 2017), or expected rewards and their relative uncertainties (Boorman et al., 2009; Badre et al., 2012). Likewise, in the case of the current study, the evidence that an ambiguous stimulus is similar to a given category must be combined with the evidence that the stimulus is dissimilar to the other possible categories. Although the dissGCM instantiates dissimilarity as a simple transformation of similarity, the involvement of rlPFC when participants place more reliance on dissimilarity-based evidence may be attributable to increasing demands for integrating evidence across several abstract representations. A decision based on pure similarity-based evidence would require no such integration. This hypothesis accords with recent findings implicating the rlPFC in evaluative processes for categorization tasks that require candidate rules to be weighed over the course of several trials, relative to matching tasks where a rule can be known with certainty following a single correct trial (Paniukov & Davis, 2018).

One question that has arisen repeatedly in the literature on the IBRE is whether it reflects an inherent irrationality in decision making. When viewed through the lens of basic similarity-based attentional processes (e.g., Medin & Edelson, 1988; Kruschke, 1996, 2001), the IBRE appears to arise from very simple learning mechanisms that are not particularly tied to higher-level rationality, and rare choices seem to indicate a lack of knowledge of the base rates. Indeed, in a separate model fit, we attempted to fit the standard similarity-based GCM to the key pattern on the ambiguous trials. However, the standard GCM was only able to predict a greater proportion of rare choices if accurate knowledge of the exemplar base-rates was eliminated (all values of *t_j_* = 1 or fit as free parameters). In contrast, accurate knowledge of the category base rates directly contributes to the greater dissimilarity-based evidence against the common category. Thus from the dissGCM perspective, participants are perfectly knowledgeable about the base rates in the present task, but they use this knowledge in a way not anticipated by pure similarity-based models. However, whether this use of dissimilarity-based evidence constitutes irrationality is a deeper question that cannot be answered based purely on the present results.

The IBRE exemplifies a case in cognitive neuroscience where independent models that predict essentially the same behavioral patterns make very different assumptions about the cognitive processes, and accordingly, brain states, involved in producing the behavior. Our findings from the test phase represent a critical step forward in an emerging area of research using multivariate fMRI to reveal that qualitatively distinct brain states may reflect the use of multiple response strategies in the face of identical stimuli (e.g., Mack et al., 2013). Consistent with past research using MVPA to decode learned selective attention (Mack et al., 2013, 2016; Leong et al., 2017; O’Bryan et al., 2018), multivoxel patterns associated with predictive features were more strongly activated than imperfectly predictive features during the learning phase. Using the same approach to decode which information participants were focusing on during ambiguous test trials, we found stronger activation of patterns associated with common compared to rare stimulus features, but importantly, this pattern only emerged in cases where participants chose the rare category. Although these results are consistent with a dissimilarity-based process where activating knowledge of the common feature provides contrastive evidence against the well-established common category, understanding the precise cognitive mechanisms that contribute to these response-dependent activation patterns remains a direction for future research. One possibility is that common choices stem from habitual response patterns that involve feature matching and require little active deliberation, and hence less activation of featural information in the associated multivoxel activation patterns, whereas more actively weighing the evidence for each category engages a “strategic guessing” process that involves ruling out the most unlikely option (Juslin, Wennerholm, & Winman, 2001).

Interestingly, while our findings argue against the prediction from similarity-based models that the IBRE arises because rare features become more similar to their associated category, the observed parameters from the model fits are consistent with a key part of similarity theory – that there is greater selective attention allocated to the rare feature dimension. Indeed, the rare feature dimensions outweighed the common features for both sets of categories in our data. However, these larger attention weights did not seem to drive greater neural similarity to the rare feature dimension in our multivoxel results. Accordingly, it is possible that our multivoxel results are not driven directly by simple feature-based attention, but instead indicate some combination of attention and memory-based retrieval of the category exemplars. This hypothesis coincides with findings from the memory literature, which have found that memory-based retrieval of non-present, associated stimuli can be detectable in activation patterns (e.g., Zeithamova, Dominick & Preston, 2012; for review, see Rissman & Wagner, 2012). Future studies may wish to combine multivoxel pattern analysis with eye-tracking (e.g., Leong et al., 2017) to better understand the unique contributions that attention and memory make to the present results.

In conclusion, using model-based fMRI analysis, we found evidence that extreme cases of base rate neglect such as the IBRE may arise from a combination of similarity-and dissimilarity-based processes. Accordingly, measures of neural activation suggest that people may be more strongly relying on evidence about how dissimilar an item is to common categories when faced with ambiguous stimuli. Further, dissimilarity processes have a unique cortical topography that includes the rostrolateral PFC, a region believed to be involved with more symbolic feature processing.

## Materials and Methods

Twenty-four healthy right-handed volunteers (age range 18 – 58; 13 women) participated in the study for $35. All protocols were approved by the Texas Tech University IRB. Two participants were excluded, one for falling asleep and the other for excessive head motion.

### Behavioral Protocol

The study consisted of three phases, localizer, learning, and test. The localizer phase consisted of two scanning runs in which participants classified images based on whether they contained a face, an object, or a scene. Each image was presented for 2.5 s during which participants were asked to respond “Scene (1), Face (2), or Object (3)?” Each trial was separated with random fixation drawn from a truncated exponential distribution with mean = 3 s. Over the duration of the localizer phase, subjects categorized 38 examples of each stimulus type. The face, object, and scene images used were black-and-white squares presented on a white background with black text. The stimuli used during the localizer runs were presented in a random order, and did not include any of the images used for the experimental task.

In the learning phase, participants learned a classic IBRE category structure based on Medin and Edelson (1988; See Figure 1). The features used for the stimuli included examples of faces, objects, and scenes not shown in the localizer phase. Participants were given an epidemiological cover story asking them to predict whether hypothetical patients would contract a disease based on the people they have been in contact with (faces), the objects they have used (objects), and the places they have been (scenes). On each trial of the learning phase, participants would see a stimulus for 3 s and were asked to answer “Disease 1, 2, 3, or 4?” This was followed by random fixation, feedback (1.75 s) in which they were told whether they were right or wrong and the correct answer, and additional fixation. The same distribution was used to generate fixations as in the localizer phase. Faces were always assigned to the imperfectly predictive feature dimensions, whereas objects and scenes were perfectly predictive and associated with only one disease (Figure 1). To ensure that no visual stimulus category differed in overall frequency, one common disease always was associated with objects and the other scenes, and likewise for rare diseases. Participants were randomly assigned to one of two conditions to balance which images were presented together during learning and test, and disease labels were randomized across participants. Within-pair stimulus position (left or right) was randomized on each trial, and the presentation order of feature pairs was randomized within each block for every participant. The learning phase was spread over three scanning runs, and four blocks of the stimulus set were presented per run, resulting in a total of12 blocks and 96 trials for the learning phase. The progression of a learning trial is depicted in the bottom panel of Figure 1.

During the test phase, participants completed trials with both new and old exemplars and classified them as “Disease 1, 2, 3, or 4?”, but no longer received feedback. New items included all possible single and two-feature combinations of the perfectly predictive features (see Table 1, Results). Trials were 3 s and separated by random fixation as described above. Like the learning phase, the test phase occurred over three consecutive scanning runs. Presentation order of the test items was randomized for each of the three runs, with participants rating two test sets per run, resulting in a total of 156 test trials.

### Image Acquisition

Imaging data were acquired on a 3.0 T Siemens Skyra MRI scanner at the Texas Tech Neuroimaging Institute. Structural images were acquired in the sagittal plane using MPRAGE whole-brain anatomical scans (TR = 1.9 s; TE = 2.44 ms; *θ* = 9°; FOV = 250 x 250 mm; matrix = 256 x 256 mm; slice thickness = 1.0 mm, slices = 192). Functional images were acquired using a single-shot T2*-weighted gradient echo EPI sequence (TR = 2.5 s; TE = 25 ms; *θ* = 75°; FOV= 192 x 192 mm; matrix = 64 x 64; slice thickness = 3 mm).

### fMRI Analysis and Preprocessing

Functional data were preprocessed and analyzed using FSL (www.fmrib.ox.ac.uk/fsl). Anatomical images were preprocessed using Freesurfer (autorecon1). Functional images were skull stripped, motion corrected, prewhitened, and high-pass filtered (cutoff: 60 s). For the model-based univariate analysis, functional images were spatially smoothed using a 6 mm FWHM Gaussian kernel. No smoothing was performed on functional data used for the multivoxel analysis. First-level statistical maps were registered to the Montreal Neurological Institute (MNI)-152 template using 6-DOF boundary-based registration to align the functional image to the Freesurfer-processed high-resolution anatomical image, and 12-DOF affine registration to the MNI-152 brain.

The model-based univariate analysis employed a standard three-level mixed effects model carried out in FSL’s FEAT program. The first-level model included an EV for stimulus presentation and two model-based parametric modulators: Similarity-and Dissimilarity-based evidence, computed from the dissGCM. Both parametric modulators were centered and scaled (z-scored) within run. Calculation of these regressors is described in the Model section. Additional explanatory variables (EVs) of no interest included motion parameters, their temporal derivatives, EVs to censor volumes exceeding a framewise displacement of 0.9mm (Siegel et al., 2014), and an EV to account for trials in which participants failed to make a behavioral response. Final statistical maps were corrected for multiple comparisons using a non-parametric cluster-mass-based correction with a cluster-forming threshold of *t* (21) = 3.52 (*p* < .001, one-tailed).

RSA was conducted using the PyMVPA toolbox (Hanke et al., 2009) and custom Python routines. In this analysis, we measured how much participants were activating scene, object, and face information on individual test phase trials by calculating mean correlation distance (1 – Pearson’s *r*) between activation patterns on each test trial and those elicited for each visual category during the localizer phase. For interpretative ease, the distances were converted to similarities using exp(-distance), and then standardized (*z*-scored) within participants. Activation patterns were estimated for each trial using a Least Squares All procedure (Mumford et al., 2012), and anatomically restricted to two ventral temporal ROIs that were maximally responsive to scene and object information in the localizer data. Specifically, pattern estimates were spatially localized in visual stimulus category-specific ROIs by creating 6-mm spheres around subjects’ peak activation within anatomically defined regions in the Harvard-Oxford Atlas associated with category selectivity (objects: left inferior posterior temporal gyrus; scenes: bilateral parahippocampal gyrus; Ishai et al., 1999; Lewis-Peacock & Postle, 2008; Lewis-Peacock et al., 2012; Grill-Spector & Weiner, 2014). The last trial of each run was automatically discarded from the multivoxel analysis to ensure stable estimation of the activation patterns for all trials. Additional explanatory variables (EVs) of no interest included motion parameters, their temporal derivatives, and EVs to censor volumes exceeding a framewise displacement of 0.9mm. Source data and scripts used to create all figures and tables (e.g., R code, PyMVPA scripts, statistical maps for the model-based fMRI analysis) are freely available online at https://osf.io/atbz7/.

## Acknowledgements

This work was supported by start-up funds to TD from Texas Tech University. The authors declare no conflicts of interest.

## References

Badre, D., & D’Esposito, M. (2007). Functional magnetic resonance imaging evidence for a hierarchical organization of the prefrontal cortex. Journal of Cognitive Neuroscience, 19, 2082–2099.

Badre, D., & D’Esposito, M. (2009). Is the rostro-caudal axis of the frontal lobe hierarchical? Nature Reviews Neuroscience, 10, 659–669.

Badre, D., Doll, B.B., Long, N.M., & Frank, M.J. (2012). Rostrolateral prefrontal cortex and individual differences in uncertainty-driven exploration. Neuron, 73, 595–607.

Boorman, E.D., Behrens, T.E.J., Woolrich, M.W., & Rushworth, M.F.S. (2009). How green is the grass on the other side? Frontopolar cortex and the evidence in favor of alternative courses of action. Neuron, 62, 733–743.

Bravata DM (2000). Making medical decisions under uncertainty. Seminars in Medical Practice, 3, 6–13.

Bunge, S.A., Wendelken, C., Badre, D., & Wagner, A.D. (2005). Analogical reasoning and prefrontal cortex: evidence for separable retrieval and integration mechanisms. Cerebral Cortex, 15, 239–249.

Bunge, S.A., Helskog, E.H., & Wendelken, C. (2009). Left, but not right, rostrolateral prefrontal cortex meets a stringent test of the relational integration hypothesis. NeuroImage, 46, 338–342.

Casscells, W., Schoenberger, A., & Graboys, T. B. (1978). Interpretation by physicians of clinical laboratory results. New England Journal of Medicine, 299, 999–1001.

Christoff, K., Prabhakaran, V., Dorfman, J., Zhao, Z., Kroger, J. K., Holyoak, K. J., & Gabrieli, J. D. (2001). Rostrolateral prefrontal cortex involvement in relational integration during reasoning. NeuroImage, 14, 1136–1149.

Davis, T., Love, B. C., & Preston, A.R. (2012a). Learning the exception to the rule: Model-based fMRI reveals specialized representations for surprising category members. Cerebral Cortex, 22, 260–273.

Davis, T., Love, B. C., & Preston, A.R. (2012b). Striatal and hippocampal entropy and recognition signals in category learning: simultaneous processes revealed by model-based fMRI. Journal of Experimental Psychology: Learning, Memory, and Cognition, 38, 821–839

Davis, T., & Poldrack, R. A. (2013). Quantifying the internal structure of categories using a neural typicality measure. Cerebral Cortex, 24, 1720–1737.

Davis, T., Goldwater, M.B., Giron, J. (2017). From concrete examples to abstract relations: The rostrolateral prefrontal cortex integrates novel examples into relational categories. Cerebral Cortex, 27, 2652–2670.

DeGutis, J., D’Esposito, M. (2007). Distinct mechanisms in visual category learning. Cognitive Affective and Behavioral Neuroscience, 7, 251–259.

De Martino, B., Fleming, S.M., Garrett, N., & Dolan, R., (2013). Confidence in value-based choice. Nature Neuroscience, 16, 105–110.

Grill-Spector, K., & Weiner, K.S. (2014). The functional architechture of ventral temporal cortex and its role in categorization. Nature Reviews Neuroscience, 15, 536–548.

Hanke, M., Halchenko, Y.O., Sederberg, P.B., Hanson, S.J., Haxby, J.V., & Pollmann, S. (2009). PyMVPA: A Python toolbox for multivariate pattern analysis of fMRI data. Neuroinformatics, 7, 37–53.

Haxby, J.V., Gobbini, M.I., Furey, M.L., Ishai, A., Schouten, J.L., & Pietrini, P. (2001). Distributed and overlapping representations of faces and objects in ventral temporal cortex. Science 293: 2425–2430.

Haxby, J.V., Connolly, A.C., & Guntupalli, J.S. (2014). Decoding neural representational spaces using multivariate pattern analysis. Annual Review of Neuroscience, 37, 435–456.

Ishai, A., Ungerleider, L.G., Martin, A., Schouten, J.L., & Haxby, J.V. (1999). Distributed representation of objects in the human ventral visual pathway. Proceedings of the National Academy of Sciences, 96, 9379.

Juslin, P., Wennerholm, P., & Winman, A. (2001). High-level reasoning and base-rate use: Do we need cue-competition to explain the inverse base-rate effect? Journal of Experimental Psychology: Learning, Memory, and Cognition, 27, 849–871.

Kriegeskorte, N., Mur, M., & Bandettini, P. (2008). Representational similarity analysis— connecting the branches of systems neuroscience. Frontiers in Systems Neuroscience, 2, 4.

Kruschke, J.K. (1996). Base rates in category learning. Journal of Experimental Psychology: Learning, Memory, and Cognition, 22, 3–26.

Kruschke, J.K. (2001). Toward a unified model of attention in associative learning. Journal of Mathematical Psychology, 45, 812–863.

Leong, Y.C., Radelescu, A., Daniel, R., & Niv, Y. (2017). Dynamic interaction between reinforcement learning and attention in multidimensional environments. Neuron, 93, 451–463.

Lewis-Peacock, J.A., & Postle, B.R. (2008). Temporary activation of long-term memory supports working memory. The Journal of Neuroscience, 28, 8765–8771.

Lewis-Peacock, J.A., Drysdale, A.T., Oberauer, K., & Postle, B.R. (2012). Neural evidence for a distinction between short-term memory and the focus of attention. Journal of Cognitive Neuroscience, 24, 61–79.

Mack, M.L., Preston, A.R., & Love, B.C. (2013). Decoding the brain’s algorithm for categorization from its neural implementation. Current Biology, 23, 1–5.

Mack, M. L., Love, B. C., & Preston, A. R. (2016). Dynamic updating of hippocampal object representations reflects new conceptual knowledge. Proceedings of the National Academy of Sciences, 113, 13203–13208.

Medin, D. L., & Edelson, S. M. (1988). Problem structure and the use of base-rate information from experience. Journal of Experimental Psychology: General, 117, 68–85.

Mumford, J. A., Turner, B. O., Ashby, F. G., & Poldrack, R. A. (2012). Deconvolving BOLD activation in event-related designs for multivoxel pattern classification analyses. NeuroImage, 59, 2636–2643.

Nosofsky, R. M. (1986). Attention, similarity, and the identification–categorization relationship. Journal of Experimental Psychology: General, 115, 39–57.

O’Bryan, S.R., Walden, E., Serra, M.J., & Davis, T. (2018). Rule activation and ventromedial prefrontal engagement support accurate stopping in self-paced learning. NeuroImage, 172, 415–426.

Paniukov, D., & Davis, T. (2018). The evaluative role of rostrolateral prefrontal cortex in rule-based category learning. NeuroImage, 166, 19–31.

Rissman, J., & Wagner, A. D. (2012). Distributed representations in memory: insights from functional brain imaging. Annual Review of Psychology, 63, 101–128.

Seger, C.A., Braunlich, K., Wehe, H.S., & Liu, Z. (2015). Generalization in category learning: The roles of representational and decisional uncertainty. The Journal of Neuroscience, 35, 8802–8812.

Siegel, J.S., Power, J.D., Dubis, J.W., Vogel, A.C., Church, J.A., Schlaggar, B.L., & Petersen, S.E. (2014). Statistical improvements in functional magnetic resonance imaging analyses produced by censoring high-motion data points. Human Brain Mapping, 35, 1981–1996.

Stewart, N., & Morin, C. (2007). Dissimilarity is used as evidence of category membership in multidimensional perceptual categorization: A test of the similarity– dissimilarity generalized context model. Quarterly Journal of Experimental Psychology, 60, 1337–1346.

Tversky, A., & Kahneman, D. (1974). Judgment under uncertainty: Heuristics and biases. Science, 185, 1124–1131.

Winman, A., Wennerholm, P., Juslin, P., & Shanks, D.R. (2005). Evidence for rule-based processes in the inverse base-rate effect. Quarterly Journal of Experimental Psychology, 58A, 789–815.

Zeithamova, D., Dominick, A.L., & Preston, A.R. (2012). Hippocampal and ventral medial prefrontal activation during retrieval-mediated learning supports novel inference. Neuron, 75, 168–179.

